# Effects of Abattoir Effluent as Biofertilizer on Pisum sativum Yield

**DOI:** 10.1101/2025.09.24.678026

**Authors:** Jighjigh Friday Gudu, Laura N Utume

## Abstract

Biological preparation containing live or latent cell of microorganisms or their metabolites, which when inoculated to seed, soil or roots of seedlings, promote plant growth and enhance harvestable yield is termed as Biofertilizers. This research was conducted to determine the growth promoting effect of some selected bacteria on pea plants. A comparative study on the effect of biofertilizers and chemical fertilizers was done on growth parameters in pea plant. An experiment was conducted in Randomized Complex Block Design (RCBD) with three replicates. The treatment were (T1=Control, T2=Biofertilizer, T3=Chemical fertilizer). Biofertilizer used in this study was abattoir effluent and chemical fertilizer was NPK fertilizers. The material consisted of two cultivars of *Pisum sativum* species. Pea seeds were planted in (21) polythene bags and left to grow to a reasonable height after which different levels of biofertilizers and chemical fertilizers were added. Biofertilizers levels of 50ml, 100ml, 150ml and chemical fertilizer of 1g, 2g and 3g were added respectively, and the procedure was repeated for twelve of the remaining polythene bags to serve as replicates while the last three polythene bags were left without fertilizer application to serve as control plant. The plants were left to develop to maturity after which growth parameters of height, number of leaves, leaf area and plant dry weight were measured for each plant as a determinant of growth. Biofertilizers had the best performance when compare with chemical fertilizers. 100ml of biofertilizers had the highest growth rate and 50ml had the least when compared to the control. Chemical fertilizers had significant difference on height, number of leaves, leaf area and however insignificant on biomass when compared to control pea plant as LSD was greater than mean difference. According to this study using biofertilizer has increased growth and growth component of pea significantly. Biofertilizers level of 100ml should therefore be adopted by farmers so as to get the maximum yield and bioconversion of organic wastes into value added organic/biofertilizers at local level should be done.

## 1. Introduction

Fertilization is one of the soil management practices which have great impact on soil and crop health. The macro and micronutrients provided via fertilizers act as adjuncts as they supplement the nutrients already present in the soil. The need of adding fertilizers to the soil is never-ending because of intensive cultivation. Fertilization of the soil can be done by adding nutrients in organic or inorganic forms. Fertilizers whether organic or inorganic, have advantages and disadvantages in context to nutrient supply, crop productivity and impact on environment (Rashid *et al*., 2016).

The use of fertilizers started when serious food shortages occurred during early 1960s. To overcome the food shortage, various technologies were introduced such as use of herbicides, pesticides and fertilizers. Fertilizers may contain one or more of the essential nutrients. Those that contain only one of the major elements are described as single, simple or straight fertilizers. Those that contain two or more of the major elements are classified as mixed or compound fertilizer. Nitrogen, phosphorus and potassium are the main plant nutrients and these three provide the basis for the major groups of fertilizers (Rashid *et al*., 2016). Although the use of fertilizers lead to the increased production of food but it had a deteriorating effect on soil health and microbial population of the soil.

The increased use of chemicals under intensive cultivation has not only contaminated the ground and surface water but has also affected the soil ecosystem (Bahadur *et al*., 2006). Contrary to the effects of chemical fertilizers on soil properties, organic fertilizers and biofertilzers improve the soil properties. Organic fertilizers are an important source of providing nutrients to plants and which avoids the use of synthetic fertilizers, pesticides, growth regulators, and livestock feed additives. Organic fertilizers include biofertilizers, animal or farm yard manure (FYM), compost, vermicompost and green manure.

Biofertilizers, a type of organic fertilizers, are emerging as an ecologically safe means of fertilization. They are defined as products containing natural occurring micro-organisms that are artificially multiplied to improve soil fertility and crop productivity (Mazid and Khan, 2014). Commonly used micro-organism as biofertilizer are *Rhizobia, Azospirillum*, Phosphate Solubilizing Bacteria (PSB), Plant Growth Promoting *Rhizobacteria* (PGPR), *Azobacter* and *Arbuscular mycorrhiza* (AM). These augment the biochemical processes in soil such as nitrogen fixation, phosphorus solubilization and mobilization, zinc solubilization, production of plant growth promoting substances and pathogen control. Biofertilizers provide an economically judicious, attractive and ecologically sound means of fertilization (Patel *et al*., 2013) and are important in maintaining sustainable agriculture. Biofertilizers not only promote plant growth and development but also reduce the cost of production as they tend to decrease the doses of chemical fertilization used. These can be used for food and fodder crops including vegetables and legume including pea plant.

*Pisum sativum* is used because of its short life span and cultivation of pea maintains soil fertility through biological nitrogen fixation while being in association with symbiotic *Rhizobium* which is present in its root nodules (Negi *et al*., 2007). Therefore, apart from meeting its own requirement of nitrogen, peas are known to leave behind residual nitrogen in soil 50-60kg/ha. Keeping all these points in view, present study was carried out to access the effect of abattoir effluent as biofertilizers on growth character of pea.

### Objectives

1. To establish a relationship between biofertilizer and pea plant growth.
2. To compare between chemical fertilizer and biofertilizer performance.

## 2. Material and Methods

### 2.1 Sample collection Seed collection

Pea (*Pisum sativum*) seed from pest infestation and diseases were collected from National Holticulture Research Institute (NIHORT), Ibadan for this research.

#### Soil collection

The top of the soil was dung and measured with a metre rule 5m below to collect loamy soil rich in nutrient for planting near the bank of River Benue. The soil sampling where done from 3-4 places and were mixed to get representative sample. 21 polythene bags were filled with uniform amount of loamy soil and weighed with weighing scale.

#### Collection and Preparation of Fertilizers

Abattoir effluent (Biofertilizer) was collected from abattoir market, Makurdi. The abattoir effluent was kept in a closed container for fourteen day, to enable the bacteria to multiply in number.

NPK fertilizers were bought from appropriate sellers in Wurukum market, Makurdi.

### 2.2 Soil Preparation and Planting

Required materials were gathered together. The polythene bags were filled with uniform amount of loamy soil and weighed with a weighing scale. Each of the polythene bags was weighed to be 3098.6 g or 3.09kg. The polythene bags were labeled “C1”, “C2”,“C3”,“B1”,“B2”,“B3”,“N1”“N2”,“N3”, so as to distinguish the various polythene bags from one another and erase confusion when applying fertilizer.

Two pea seeds were planted in each polythene bag about half way down. The peas were watered in each polythene bag so that the top is moist. The peas were watered daily until the seedlings have germinated to a significant height.

### 2.3 Field Management

Watering was carried out twice a day, morning and evening. Weeding commenced at 2 weeks after sowing of pea seed and subsequent weeding was carried out as at when due.

### 2.4 Fertization

The abattoir effluent was stir gently and a measuring cylinder was used to measure the biofertilizer into 3 different levels, that is 50ml, 100ml and 150ml. Small holes were made beneath the root zone at a considerable distance away from the pea plant and it was poured into the holes and covered with little soil.

A weighing balance was used to weighed NPK fertilizers at 1g, 2g and 3g. They were also poured into the small holes and covered with little soil.

### 2.5 Measurement of Grwoth Parameters

After the pea plants have grown sufficiently and developed firm leaves, and other structures, measurement were taken from all experimental set up.

Plant height of pea plant was recorded from ground to base of last fully opened leaf with the help of a scale. The mean values for height were expressed in centimeters (cm)

Number of leaves per plant where determined by counting and were recorded at different time intervals viz. 25, 35, and 45 days.

To obtained the biomass, Plants where cut from root-shoot junction to get root and shoot weight. The roots were washed with tap water twice or thrice to remove all the dust particles. The plants were air dried first to take the fresh weight and then dried in an oven at 60-70C days they were dried to constant weight. Dry weight was recorded using electronic weighing balance and expressed in grams (g).

Leaf area was measured by placing the ruler from the base of the leaf then across.

### 2.6 Statistical Analysis

Data generated was analyzed using analysis of variance (ANOVA) statistical test to describe the relationship between biofertilizers and pea plant growth. Least significant difference (LSD) also used to compare the means. The confidence level was placed at 5%.

## 3. Results

Table 1 shows that, the biofertilizer application at all rates resulted in a significant difference and chemical fertilizer of 1g had no significant difference.

**Table 1:** Effect of fertilizer on Plant Height (cm)

| **Treatment** | **Mean Plant Height (cm)** | **Difference from control** |
| --- | --- | --- |
| **Bio fertilizer (50ml)** | 15.467 | 6.801<sup>*</sup> |
| **Bio fertilizer (100ml)** | 19.333 | 10.667<sup>*</sup> |
| **Bio fertilizer (150ml)** | 18.333 | 9.667<sup>*</sup> |
| **NPK (1g)** | 13.967 | 5.301<sup>ns</sup> |
| **NPK (2g)** | 17.733 | 9.067<sup>*</sup> |
| **NPK (3g)** | 17.333 | 8.667<sup>*</sup> |
| **Control** | 8.666 | - |

Figure 1 show that biofertilizer had the highest significant difference with the mean difference of (9.045) as compared to NPK fertilizers with (7.678).

**Figure 1:**
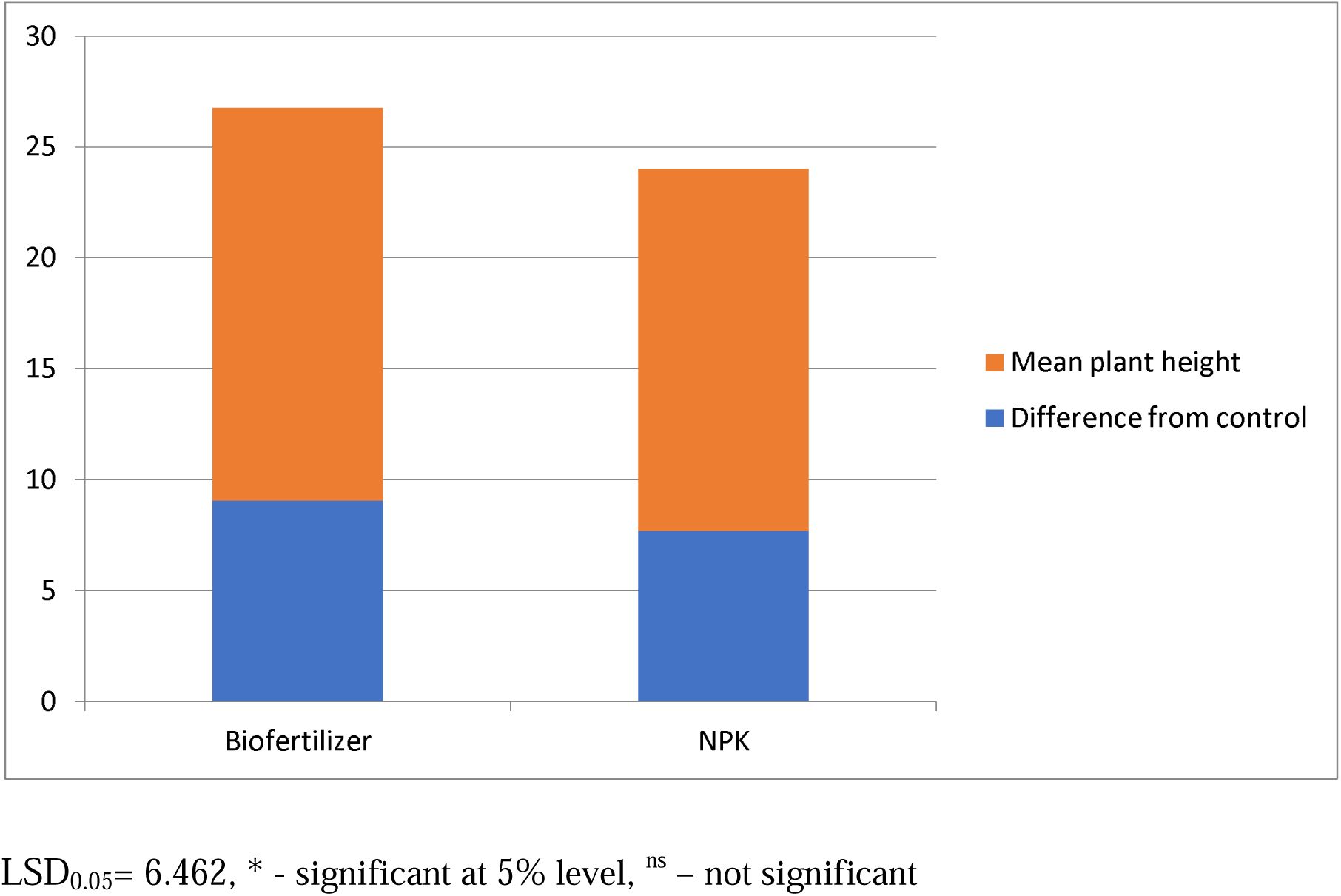
Effect of fertilizer on plant height. LSD_0.05_= 6.462, * - significant at 5% level, ^ns^ – not significant

Table 2 show that both biofertilizer and chemical fertilizer application at all rates resulted in a significant difference

**Table 2:**
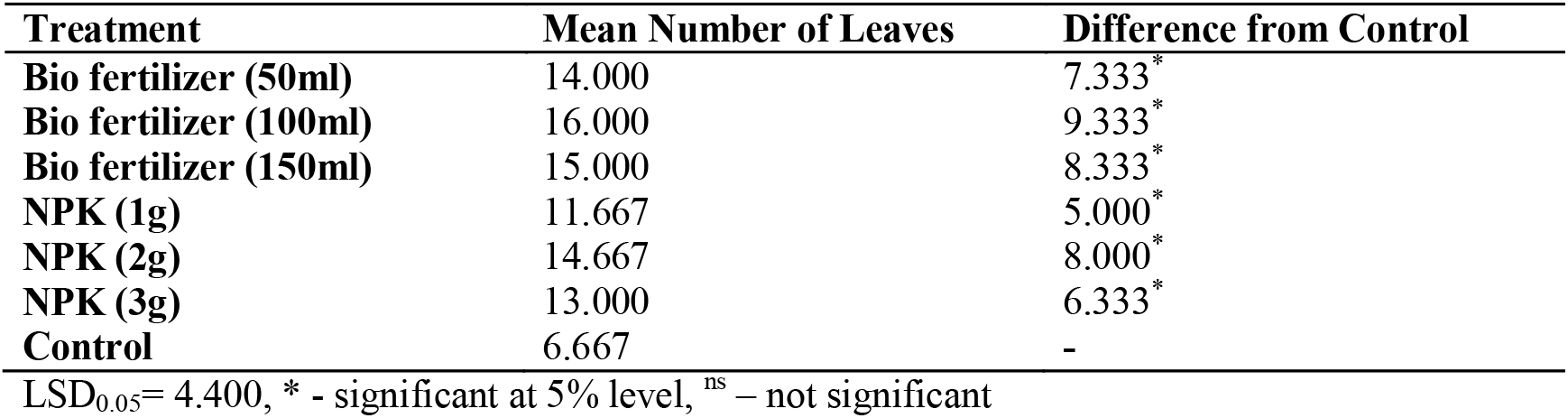
Relationship between fertilizer and leaf production.

| **Treatment** | **Mean Number of Leaves** | **Difference from Control** |
| --- | --- | --- |
| **Bio fertilizer (50ml)** | 14.000 | 7.333<sup>*</sup> |
| **Bio fertilizer (100ml)** | 16.000 | 9.333<sup>*</sup> |
| **Bio fertilizer (150ml)** | 15.000 | 8.333<sup>*</sup> |
| **NPK (1g)** | 11.667 | 5.000<sup>*</sup> |
| **NPK (2g)** | 14.667 | 8.000<sup>*</sup> |
| **NPK (3g)** | 13.000 | 6.333<sup>*</sup> |
| **Control** | 6.667 | - |

Figure 2 show that biofertilizer had the highest significant difference with the mean difference of (8.3333) as compared with NPK fertilizer with the mean difference of (6.444).

**Figure 2:**
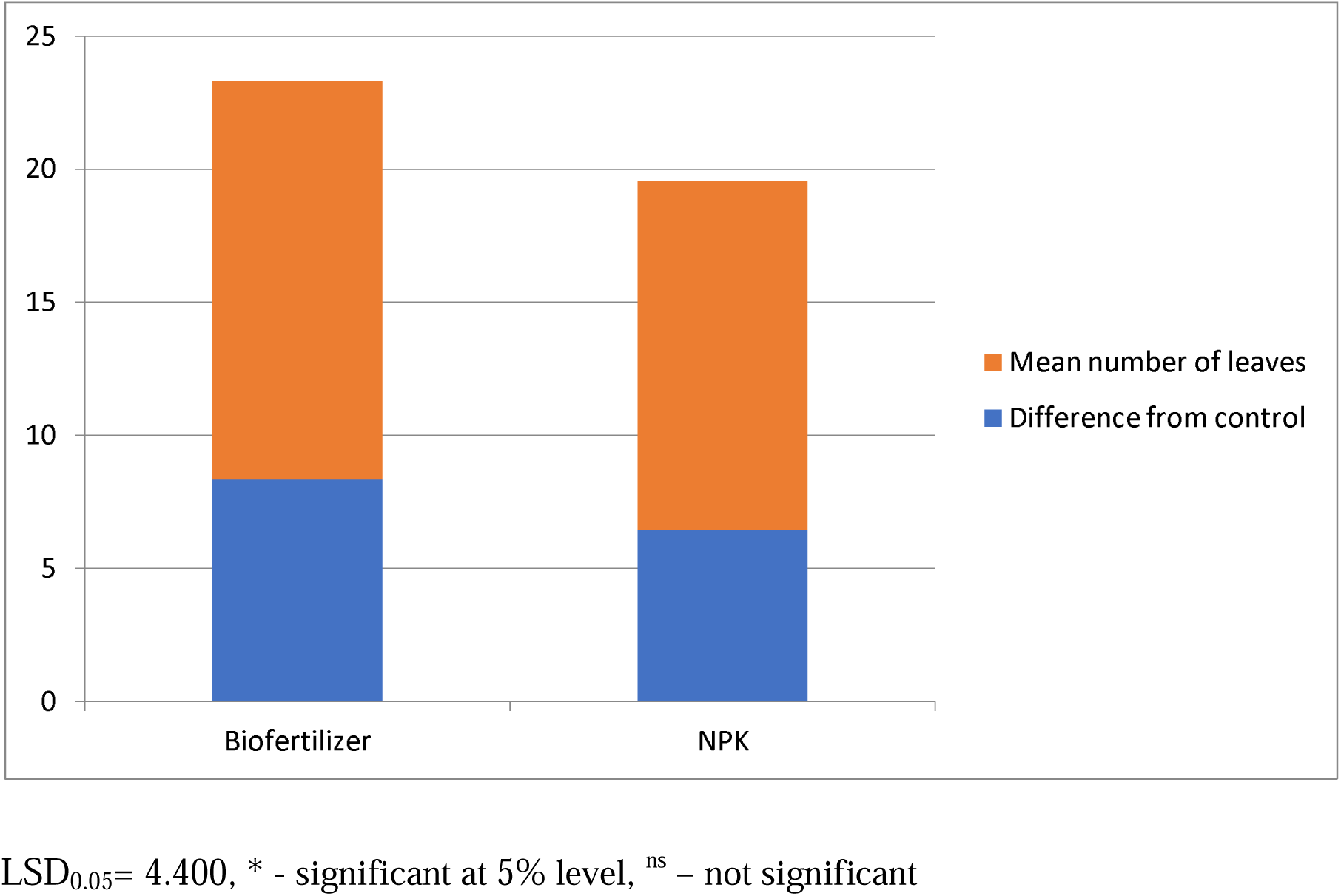
Relationship between fertilizers and leaf production. LSD_0.05_= 4.400, * - significant at 5% level, ^ns^ – not significant

Table 3 shows that 50ml of biofertilizer was not significant and the rest were significant whereas 1g of chemical fertilizer was not significant and the rest were significant.

**Table 3:** Effect of fertilizer on leaf area (cm^3^)

| **Treatment** | **Mean Leaf Area (cm<sup>3</sup>)** | **Difference from Control** |
| --- | --- | --- |
| **Bio fertilizer (50ml)** | 2.200 | 0.633<sup>ns</sup> |
| **Bio fertilizer (100ml)** | 2.967 | 1.400<sup>*</sup> |
| **Bio fertilizer (150ml)** | 2.833 | 1.266<sup>*</sup> |
| **NPK (1g)** | 2.033 | 0.466<sup>ns</sup> |
| **NPK (2g)** | 2.633 | 1.066<sup>*</sup> |
| **NPK (3g)** | 2.467 | 0.900<sup>*</sup> |
| **Control** | 1.567 | - |

Figure 3 show that biofertilizer had the highest significant difference with the mean difference of (1.100) as compared to chemical fertilizer with the mean difference of (0.811).

**Figure 3:**
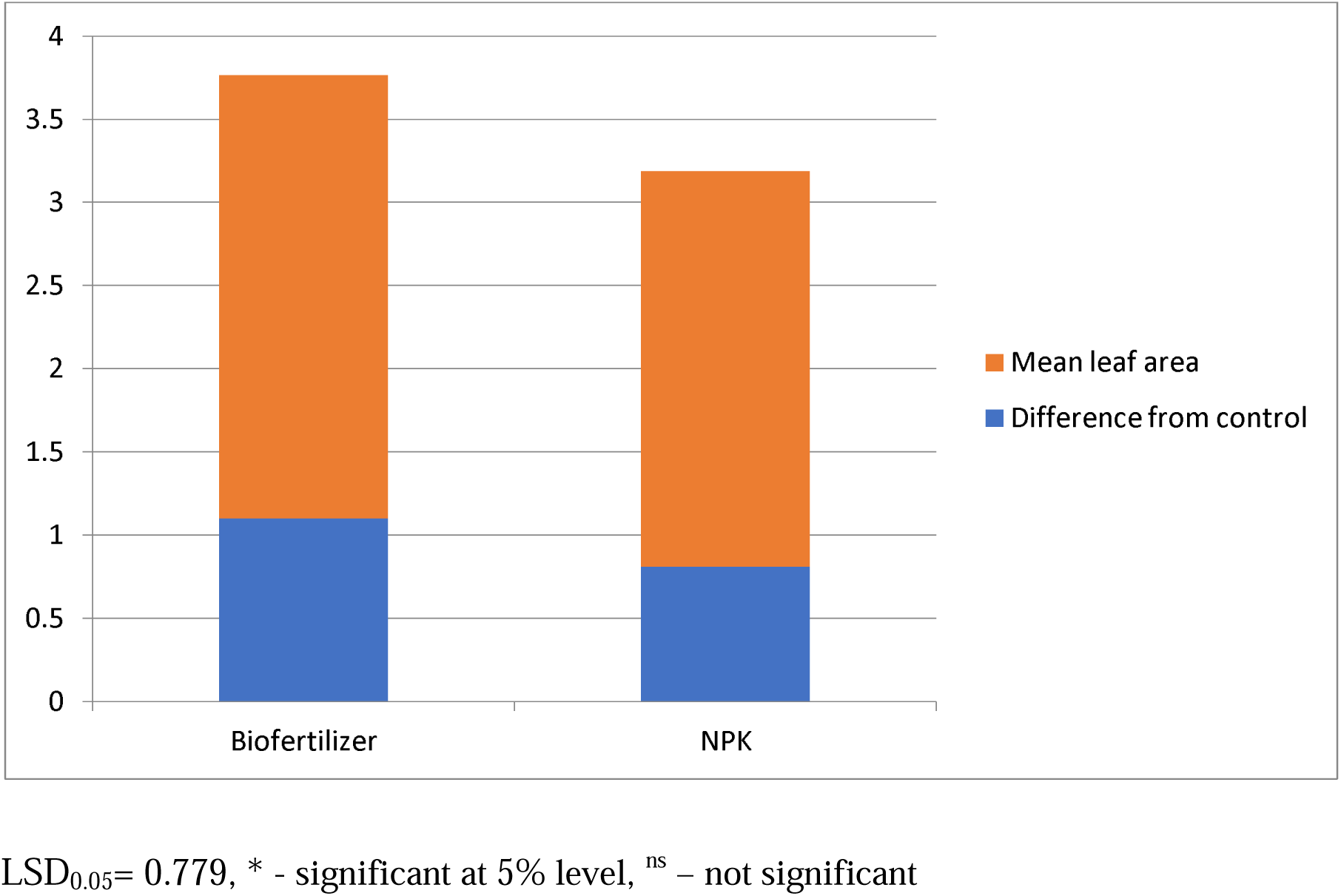
Effect of fertilizer on leaf area (cm^3^) LSD_0.05_= 0.779, * - significant at 5% level, ^ns^ – not significant

Table 4 showed that 50ml of biofertilizer was not significant and 100ml, 150ml were significant whereas none of the rates of chemical fertilizer on biomass.

**Table 4:** Relationship between fertilizers and Biomass.

| **Treatment** | **Mean Diameter** | **Difference from Control** |
| --- | --- | --- |
| **Bio fertilizer (50ml)** | 0.547 | 0.050<sup>ns</sup> |
| **Bio fertilizer (100ml)** | 1.000 | 0.503<sup>*</sup> |
| **Bio fertilizer (150ml)** | 0.687 | 0.190<sup>*</sup> |
| **NPK (1g)** | 0.480 | -0.017<sup>ns</sup> |
| **NPK (2g)** | 0.553 | 0.056<sup>ns</sup> |
| **NPK (3g)** | 0.537 | 0.040<sup>ns</sup> |
| **Control** | 0.497 |  |

Figure 4 showed that biofertilizer was significant with the mean difference of (0.247) while chemical fertilizer was not significant on biomass.

**Figure 4:**
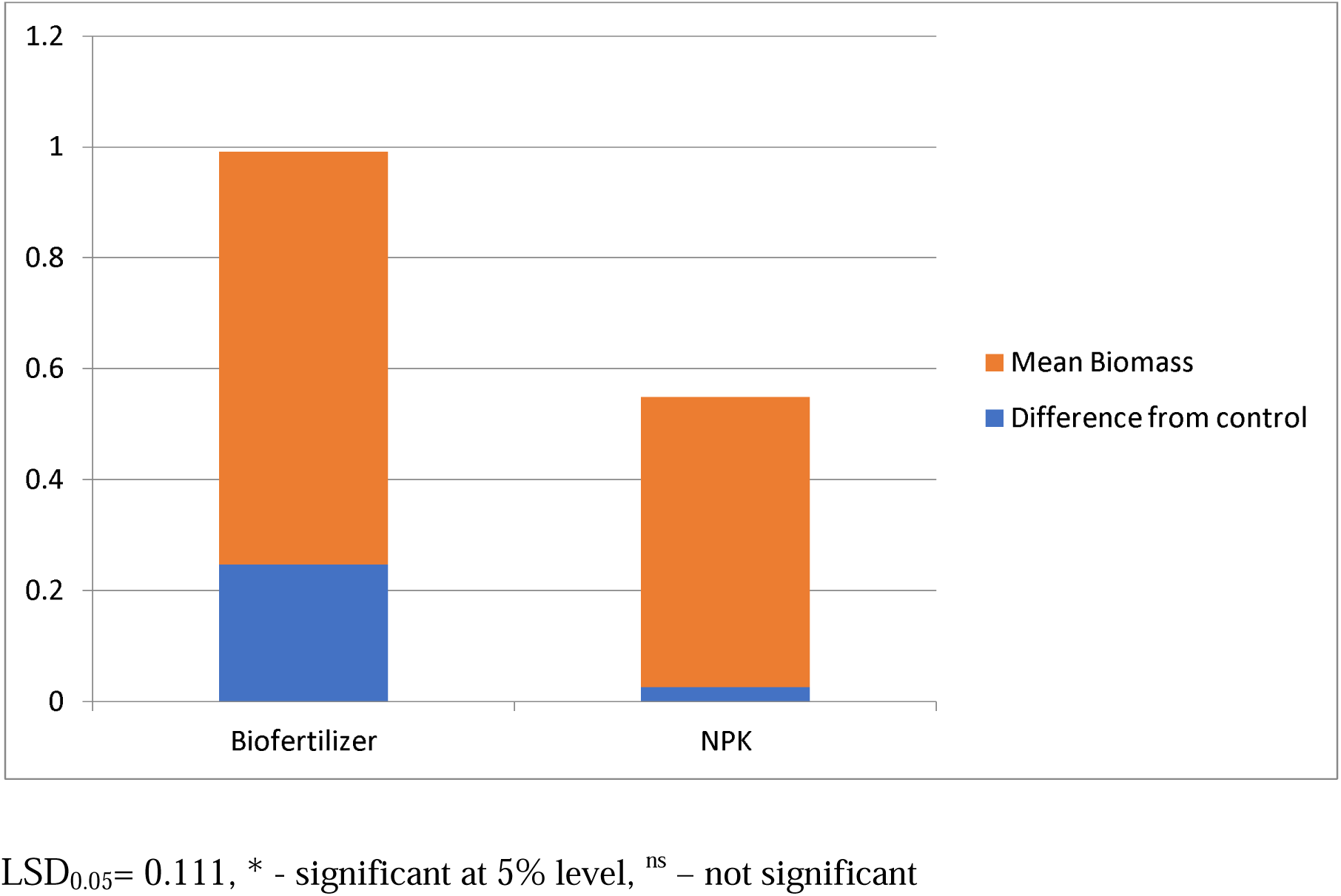
Relationship between fertilizer and biomass. LSD_0.05_= 0.111, * - significant at 5% level, ^ns^ – not significant

## 4. Discussion

Growth in plants (peas) is usually measured in terms of different parameters which may include the height of the plant, the number of leaves, leaf area and the biomass (plant dry weight) produced by the plant etc. Increase or decrease in growth is therefore measured in terms of these parameters so as to adjust conditions to favour growth of plants. In this study, both biofertilizers and NPK fertilizers had a significant effect on the growth of pea plants (*pisum sativum*). However, the pea plants treated with biofertilizer had the best performance as compared to the chemical fertilizer, and these results are in agreement with the work done by Bahadur et al., 2006. The result depicted that, 100ml of biofertilizer gave the highest growth rate. This concurs with a study done by Kurma *et al*., 2015, they reported that 100% NPK + vermicompost + biofertilizers gave the maximum growth was significant over all other treatments having single fertilizer or none.

Application of biofertilizers and NPK fertilizers had significant effect on the height of pea plants. Figure 1, show that biofertilizer had the highest significant difference with the mean difference of (9.045) as compared to NPK fertilizer with mean different of (7.678). Similarly, Dileep *et al*., 2001 produced this result when they work with *Rhizobium* and phosphate solubilizing bacteria, that they performed their study on the fennel plant. An experiment was also conducted by (Singh, 2012) on pea plant and the result shows that single inoculation either with Rhizobium or PSB and dual inoculation with both of them had significant effect on plant height which has the ability to convert nutritionally important element from unavailable to available form through biological process.

Similar trend was observed in number of leaves, and figure 2 shows that biofertilizer also had highest significant difference with a mean difference of (8.333) as compared with NPK fertilizer with the mean difference of (6.444). Increasing the number of leaves per plant could be caused by increase in plant height that was the result of improved nutrient absorption of phosphorus and nitrogen. Nitrogen as part of the protein compounds, enzymes, effective compounds in energy transfer, takes part in structure of DNA, present in the structure of chlorophyll and has a direct impact on vegetative growth. These results are in accordance with the finding of ( Patel *et al*., 1996). The result also agrees with finding of (Shanmugan and Veraputhran, 2000) that application of green manure and biofertilizer stimulated the growth of plant of plants with more number of tillers and growth of plants with more number of tillers and broader leaves in rice.

Figure 4 show that biofertilizer had significant effect on plant dry weight (biomass) whereas NPK fertilizer had no significant effect on (biomass) plant dry weight. The results are in accordance with Kandil *et al*., 2004, their experimental plant was however sugar beet. To justify these results, it could be suggested that phosphate solubilizing bacteria in these fertilizer levels had a positive impact on plant growth and increased plant dry weight. This can be caused by stimulating secretion of growth hormones which is produced by this bacteria and their effect on plant growth. They also reported that, the use of biological fertilizers in sugar beet, significantly increased plant dry weight.

## 5. Conclusion

The findings of this study showed that, pea growth parameters such as plant height, number of leaves, leaf area, leaf index, number of branches, biomass increased significantly with increased in biofertilizers level and NPK fertilizer level. So the impact of biofertilizer on examined traits was more than NPK fertilizer. Overall, the results obtained in this experiment showed that, between examined fertilizer levels, 100ml treatment of Biofertilizers provided the best conditions for achieving maximum growth in pea plant.

## Recommendations

The comprehensive development and utilization of biofertilizers is necessary, this study recommends that:

1. Necessary legislation for monitoring biofertilizers, their quality and hazardous effects, if any, on plants and human beings.
2. Government may subsidize or provide loans for small scale production units of biofertilizers at local level.
3. There is dire need of culture collections (CC)/gene bank of microorganisms in the country.
4. Research focus more on soil microbiomes, soil metagenomics, transcriptomics and key genes of active microorganisms associated with nodulation, growth regulators, disease suppressing and nutrient cycling in soil.

